# The Impact of Visual Perturbations on Balance Control during Walking

**DOI:** 10.1101/2025.04.29.651186

**Authors:** Yaqi Li, Eugenie Lambrecht, Sjoerd M. Bruijn, Jaap H. van Dieën

## Abstract

Visual perturbations may lead to an illusory self-motion and affect balance control. We studied effects of different visual perturbations in 16 healthy young participants walking on a treadmill, by assessing foot placement and center of mass (CoM) states. Three different visual perturbations were applied: fixating on a stationary target while the background moved to the right (MB), tracking the target moving rightward over a stationary background with head rotation (MT-HR), and tracking the moving target with eye movement only (MT-EM). Deviations of foot placement, CoM and trunk orientation due to the visual perturbation were assessed. Linear models were fit to the kinematic data to predict foot placement from CoM state at mid-swing. Over the whole trial, MT-HR and MT-EM caused an increase in step width variability, CoM position variability and RMS of foot placement errors simultaneously. During visual perturbation epochs specifically, in MB, a left deviation of foot and CoM trajectories was observed from the start of the background movement. In MT-HR and MT-EM, a right deviation of foot and CoM trajectories was observed only after the target had stopped moving. Contrary to our expectations, foot placement errors did not coincide with subsequent CoM deviations in the opposite direction. An obvious change in frontal plane trunk orientation was found only in MT-HR. While all visual perturbations affected control of the CoM trajectory in the frontal plane, these effects appeared caused by effects on control of heading as well. Head rotation appears to additionally disturb balance through a coupling with trunk orientation.

**Summary statement:** Visual perturbations during walking alter foot placement and body motion in humans, revealing how eye and head movements influence walking balance.

## Introduction

Balance control is a dynamic process that stabilizes the trajectory of the body’s center of mass (CoM) against the destabilizing effects of gravity encountered by humans in upright postures (Bruijn and van Dieen, 2018). The main mechanism to maintain balance during walking is the selection of the appropriate location for foot placement in relation to the ongoing movement of the CoM. This control has been identified by a linear relation between the CoM state during the swing phase. This coordination of foot placement to CoM state has been found in locomotion of flies, mice, and humans (De Comite and Seethapathi, 2025 preprint), and is thought to be achieved at least in part actively (van Leeuwen, Bruijn et al. 2022). For this control, the state of the CoM is estimated from sensory information from the visual, vestibular, and proprioceptive systems (Perry et al., 2001; Betker et al., 2008; Magnani et al., 2023; Li et al., 2025). This information is integrated by the central nervous system (CNS), which then sends signals to alpha-motor neurons to execute muscular responses to achieve the appropriate foot placement.

The visual system plays a fundamental role in navigation and predation in animals (Barron and Srinivasan, 2006; Bhagavatula et al., 2011). When encountering objects of interest, animals typically fixate or track them using eye or head movements. However, such behaviors during locomotion can disrupt the use of optical information for maintaining balance. For human beings, fixating on a stationary target while the background is moving is known to perturb balance control. Previous studies have shown that such background movement can lead to an illusion of self-motion in the direction opposite to the background movement (Nakamura, 1996; Nakamura et al., 2016), triggering balance responses that move the CoM in the direction of the background movement. In walking, it has been found that people step in the direction of the illusory self-movement and displace their CoM in the opposite direction (Reimann et al., 2018).

Another potential visual perturbation during walking may be caused by visual tracking of moving targets. Tracking a moving target causes relative movement of the visual background on the retina, which may induce a similar illusion as described above. Tracking a moving target is usually completed by coordinated eye and head rotation movements (Gadotti et al., 2016; Srivastava et al., 2018; Yamazoe et al., 2019). Some situations, such as when a large field of view shift is needed or when a target moves too fast for visual pursuit, require concomitant head movement (Rinaudo et al., 2019). These head movements affect vestibular and neck proprioceptive information and may have possible mechanical effects on balance. Previous literature has shown changes in the gait pattern due to visual tracking and related head movements. Magnani et al (2020) demonstrated that yaw head motion made gait less stable. Vallis and Patla (2004) showed that head rotation during walking coincided with a mediolateral deviation of the CoM trajectory.

As stated above, the CoM trajectory during walking is mainly stabilized by foot placement (Bruijn and van Dieen, 2018; van Leeuwen et al., 2022). However, it is not yet clear how fixating on a stationary target over a moving background and tracking a moving target affect control of the CoM by foot placement. For the latter condition, it has not been studied how rotating the head and re-directing the gaze contribute to effect of visual perturbations. In the current study, we investigated deviations in CoM trajectories and foot placement during the aforementioned visual perturbations in walking humans. We identified the linear relation between foot placement and preceding CoM state, in order to assess how steps taken during the visual perturbations contributed to balance regulation. During normal walking, these foot placement errors vary around zero. During the perturbations, however, we expected systematic deviations from zero. We hypothesized that (1) the foot placement error will increase and (2) people will step in the direction opposite to the background movement and in the direction of the target movement during these visual perturbations, which will subsequently cause displacement of the CoM in the opposite direction.

## Methods

### Participants

Sixteen healthy individuals (age: 23.4±3.9 years, height: 1.70±0.09 m, weight: 62.2±11.7 kg) volunteered for this study. All participants reported themselves to be free from any neurological or musculoskeletal disorders that could negatively affect their balance or walking performance. All participants walked without assistive devices. Ethical approval was obtained from the ethics committee of the Faculty of Behavioral and Movement Sciences at Vrije Universiteit Amsterdam (VCWE-2023-170). All procedures conformed to the Declaration of Helsinki and participants signed informed consent before participation.

### Instruments

All measurements were conducted on a system that integrated a motion capture system sampling at 50 samples/sec (Optotrak, Northern Digital, Waterloo ON, Canada) with an instrumented treadmill (Motek-Forcelink, Amsterdam, Netherlands), and a data projector. Clusters of three markers were affixed to the participant’s bodies at the following locations: the posterior surfaces of both heels, the sacrum at the level of S2, the trunk at the level of T6, and the back of head. A screen (2.50 m* 1.56 m) was placed 1.60 m in front of the participants with the projector behind it to present the visual perturbations. **Fig. 1A** shows the full set-up.

**Figure 1.**
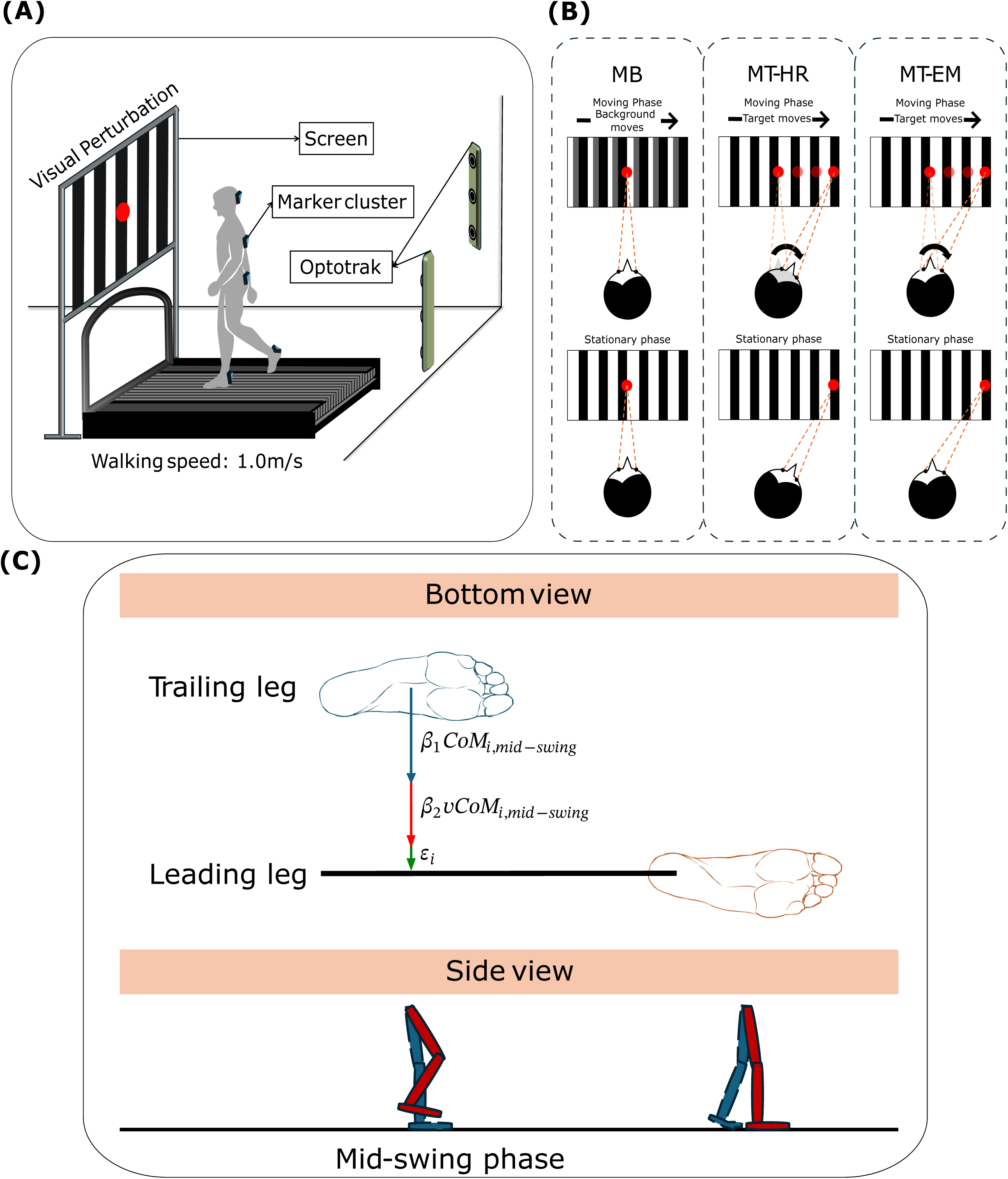
Illustration of the experimental setup (A), visual perturbation conditions (B) and foot placement model (C).

### Visual perturbations

A background of 12 uniformly distributed black-and-white vertical stripes (0.205 m * 1.560 m) was presented on the screen. A red target dot measuring 12 cm in diameter was positioned at the center of the screen. The height of the red target was adjusted to be at the participant’s eye level. Participants walked under four different conditions: (a) Normal walking (NW), in which the whole scene was stationary and participants were asked to walk normally. (b) Moving background (MB), during this perturbation, the red target dot was fixed in the middle, while the black-and-white background moved horizontally from the middle to the right side corresponding to 45 degrees in 4s, stayed at the side for 8s, then it went back to the middle. In this condition, participants were asked to keep looking at the stationary red target dot in the middle. (c) Moving target with head rotation (MT-HR), during this perturbation, the black-and-white background was stationary, while the red target dot moved horizontally to 45 degrees from the center to the right side in 4s and stayed at the side for 8s, then moved back to the middle. Participants were asked to track the moving target. (d) Moving target with eye movement (MT-EM), during this perturbation, the scene was the same as during MT-HR, but the participants were asked to track the horizontally moving target with their eyes only while keeping their head stationary. All conditions are illustrated in **Fig. 1B**.

Visual perturbations were triggered at right heel strikes. Approximately 14 perturbations in total were provided in each walking trial. The first perturbation was triggered at the 20th right heel strike. Six to eight heel strikes occurred randomly after every perturbation before a new perturbation was provided.

### Procedures

The treadmill speed was set to 1.0m/s. To enhance immersion, the lights in the lab were turned off, and the curtains were closed. Before the actual measurement, there was a 10-minute period during which participants were familiarized with the walking speed and all perturbations. Subsequently, a normal walking trial was conducted, followed by the three perturbation trials in a randomized order. Each trial lasted 5 minutes. A 5-minute seated rest was given between trials.

### Data analysis

The mediolateral CoM position was approximated as the average position of the pelvis marker cluster. Mediolateral foot positions were represented by both heel clusters. Trunk orientation in the frontal plane was calculated based on the trunk cluster relative to the laboratory coordinate system. Heel strikes were determined from the maximum values of the heel markers’ position in the anteroposterior direction.

We calculated gait characteristics which reflect balance over the whole walking trial, including step width, step width variability, step frequency, CoM position variability, trunk orientation variability, and RMS of foot placement errors. Variability of the kinematic variables was quantified as the standard deviation over steps (step width) or the standard deviation averaged over phases of the time-normalized stride (CoM position and trunk orientation). Step width was defined as the mediolateral distance between the heel positions of both feet at heel strike. Step frequency was defined as the number of steps per minute. The RMS of foot placement errors was defined as the RMS error between actual foot placement and the foot placement predicted by the model described below. Variables are presented as mean ± standard deviation. Variables that did not follow a normal distribution are presented as median (interquartile range, IQR).

The continuous variables were low-pass filtered at 0.25Hz (2nd order bidirectional Butterworth filter) to eliminate the fluctuations related to the stride cycle. We analysed the time series of foot position, foot placement error, CoM position, and trunk orientation in detail from the start of the visual perturbation to its end and divided these episodes into two phases: moving and stationary phases (**Fig. 1B**). For all variables during the visual perturbation epochs, we subtracted the value at the first sample of the epoch. Peak position during each phase was selected as the maximum absolute value. We determined the average values of these variables among subjects over perturbations. Foot placement errors in both phases of the visual perturbation were extracted from the foot placement model described below.

We applied a foot placement model to show step by step foot placement regulation during visual perturbations. We used the complete time series of each trial to determine a linear model predicting foot placement by the preceding CoM state. The model linked the variance in mediolateral foot placement at heel strike to the variance in mediolateral CoM position and velocity during the preceding swing phase (Wang and Srinivasan, 2014; Arvin et al., 2018; van Leeuwen et al., 2020) as:

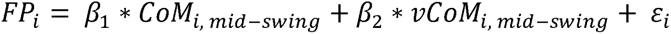

with foot placement being foot placement of the leading leg (*FP*) at step *i* and *CoM_i, mid-swing_* being the position of the CoM at mid-swing relative to the trailing foot’s position on the ground, and *vCoM_i, mid-swing_* referring to the velocity of the CoM at mid-swing (**Fig. 1C**). *β*_1_ and *β*_2_ are regression coefficients and *E_i_* is the error for every step. The RMS of *ε* over all steps was defined as the residual error in foot placement.

For each participant, we counted the number of steps from the initial right heel strike to the end of the movement and stationary phases. We used the minimum number of steps per phase over all participants to analyse foot placement errors. In the stationary phase, we selected the first right step as step 1. Specifically, foot placement errors were analysed for the first seven steps in the moving phase and the first eleven steps in the stationary phase.

### Statistics

All statistical analyses were performed in Matlab (R2023a, The Mathworks Inc., Natick, MA). For all statistics tests, we checked the assumption of normality using a Shapiro-Wilks test. To assess the overall effects of visual perturbations on gait characteristics, we used a repeated measures analyses of variance with Visual perturbation as a factor. For variables that did not follow a normal distribution (i.e. CoM variability and RMS of foot placement errors), we used a Friedman test. For the perturbation epochs, we used a one-sample *t*-test to assess if averaged peak foot positions, peak CoM positions, peak trunk orientations, and foot placement errors per step were significantly different from zero. For steps where the foot placement error that didn’t follow a normal distribution, we used Wilcoxon signed-rank test. We conducted repeated measures analyses of variance with two factors (Visual perturbation and Phase) for peak foot positions, peak CoM positions, peak trunk orientations in the perturbation epochs (MB, MT-HR and MT-EM). If we found main effects or interaction effects to be significant, we performed Bonferroni corrected pairwise post-hoc comparisons to test for differences among visual perturbations for each phase (moving and stationary phases). We calculated the effect size to provide an estimate of the magnitude of observed effect: *d* (Cohen’s d) for one-sample t-tests, *η*^2^ (Eta squared) for ANOVA, *W* (Kendall’s W) for Friedman test, and *r_rb_* (rank-biserial correlation) for Wilcoxon signed-rank test. For all repeated measures analyses, if the assumption of sphericity was violated, the Greenhouse-Geisser correction was be applied to adjust the degrees of freedom. For all analyses, the significance level (p-value) was set at 0.05.

## Results

### Destabilizing effects of visual perturbations over the whole trial

There were no significant effects of Condition on step width and step frequency. Step width variability was significantly affected by Visual perturbation (F_visual perturbation_ (df = 3,45) = 10.744, p < 0.001, *η*^2^= 0.417) (**Fig. 2**). Step width variability in the NW condition (0.022 ± 0.005m) was significantly lower than in the MT-HR (0.026 ± 0.005m) and MT-EM (0.027 ± 0.005m) conditions, corresponding to 84.6% and 81.4% of the MT-HR and MT-EM values, respectively (*p* < 0.01, *d* = -0.795; *p* < 0.001, *d* = -1.079). There was no significant difference between MB and the other conditions. CoM position variability was significantly affected by Visual perturbation (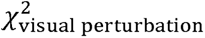(df = 3,45) = 30.225, p < 0.001, *W* = 0.630). CoM position variability in the NW condition was significantly smaller than in the MB, MT-HR and MT-EM conditions (p < 0.001, *r_rb_* = -0.897; p < 0.001, *r_rb_* = -1.000; p < 0.001, *r_rb_* = -1.000, respectively). CoM position variability in NW (0.013 (0.012-0.015) m) was 68.4%, 46.4% and 41.9% of the MB (0.019 (0.018-0.021) m), MT-HR (0.028 (0.022-0.039) m) and MT-EM (0.031 (0.025-0.034) m) values. CoM position variability in the MB condition was significantly smaller than MT-HR and MT-EM conditions (p < 0.01, *r_rb_* = -0.662; p = 0.012, *r_rb_* = -0.676), corresponding to 67.9% and 61.3% of the MT-HR and MT-EM values. MT-HR and MT-EM were not significantly different from each other (p = 0.829, *r_rb_* = -0.235).

**Figure 2.**
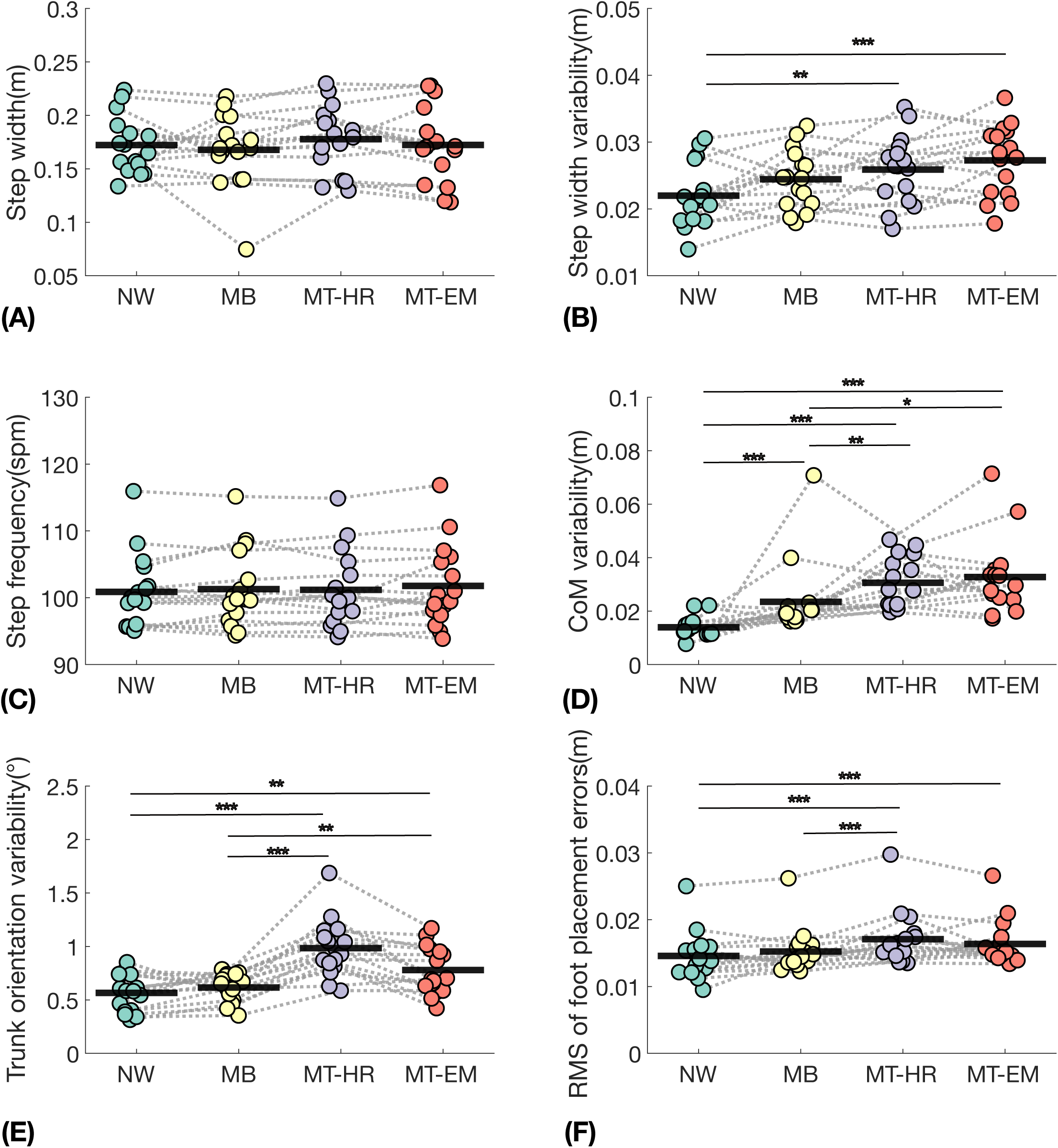
Effects of Visual perturbation conditions on step width (A), step width variability (B), step frequency (C), CoM position variability (D), trunk orientation variability (E), and RMS of foot placement errors (F). Variability of the kinematic variables was quantified as the standard deviation over steps (mean ± s.d., n =16). The black thick lines represent the mean value across individuals. Repeated measures analyses of variance were used for statistical analysis. For the variables that did not follow a normal distribution including CoM variability and RMS of foot placement errors, Friedman test was performed. Significant effects in the post-hoc tests are indicated by asterisks (* denotes p < 0.05, ** denotes p < 0.01, *** denotes p < 0.001). NW: normal walking, MB: moving background, MT-HR: moving target with head rotation, MT-EM: moving target with eye movement. Data point represents the mean values over the time series for each individual.

Trunk orientation variability was significantly affected by Visual perturbation (F _visual perturbation_ (df = 3,45) = 31.534, p < 0.001, *η*^2^ = 0.678). During NW, it was significantly smaller than during MT-HR (p < 0.001, *d* = -2.044) and MT-EM (p < 0.01, *d* = -1.044) conditions. Trunk orientation variability in NW (0.566 ± 0.172°) was 57.5% and 72.6% of the MT-HR (0.984 ± 0.267°) and MT-EM (0.780 ± 0.223°) values. MT-HR and MT-EM conditions also had significantly greater trunk orientation variability than MB (p < 0.001, *d* = 1.799; p < 0.01, *d* = 0.800). Trunk orientation variability in the MB condition was not significantly different from that in NW (p = 0.196, *d* = 0.245).

The RMS of foot placement errors was significantly affected by Visual perturbation (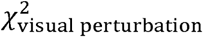 (df = 3,45) = 29.775, p < 0.001, *W* = 0.620). It was significantly smaller in NW (0.014 (0.013-0.016) m) than in MT-HR (0.016 (0.015-0.018) m) and MT-EM (0.015 (0.015-0.017) m), corresponding to 87.5% and 93.3% of the MT-HR and MT-EM values, respectively (*p* < 0.001, *r_rb_* = -0.926; *p* < 0.001, *r_rb_* = -0.971). RMS of foot placement errors in MT-HR was significantly greater than in MB (p < 0.001, *r_rb_* = 0.956), while MB was not significantly different from NW (p = 0.834, *r_rb_* = 0.221).

### Destabilizing effects during the visual perturbation

Qualitatively, foot trajectories changed during the visual perturbations (**Fig. 3**). In the MB condition, participants placed their feet in the direction opposite to the background movement from the start of the moving phase. A peak was reached at the beginning of the stationary phase and then the foot trajectories tended to gradually return to the original location. In the MT-HR condition, participants placed their feet more in the movement direction of the target during the moving phase and continued deviating throughout the stationary phase. In the MT-EM condition, participants placed their feet more in the direction of the target from the end of the moving phase and continued deviating throughout the stationary phase.

**Figure 3.**
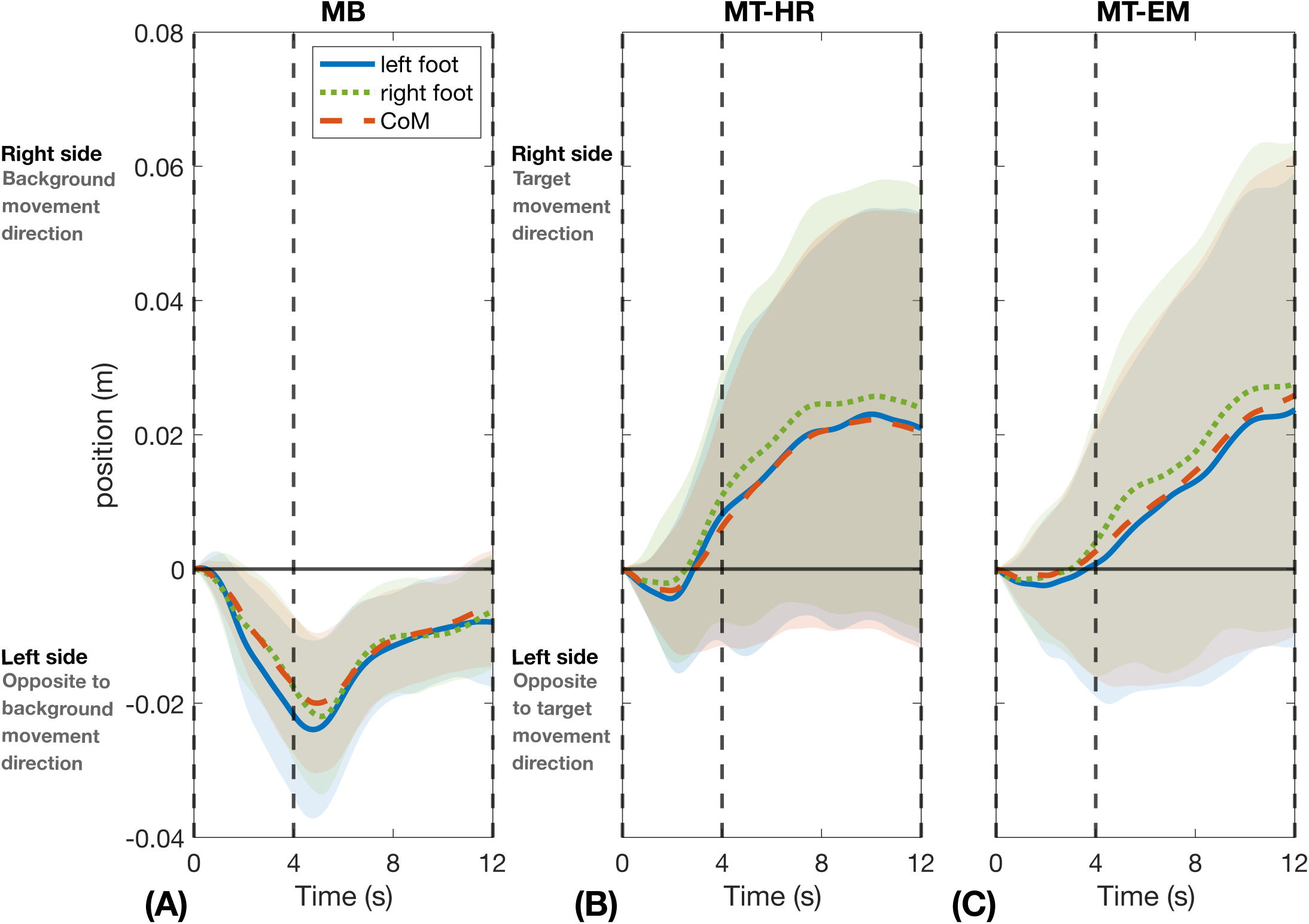
Deviations in foot and CoM positions during visual perturbation epochs. Data represent mean value across repeated visual perturbations and participants and the shaded area represents the between-participants standard deviations (mean ± s.d., n =16). All the data were referenced to the first sample of the visual perturbation epoch. From left to right: (A) MB: moving background. (B) MT-HR: moving target with head rotation, (C) MT-EM: moving target with eye movement. The vertical dashed lines separate the different phases: the line at 0s marks the start of the moving phase, the line at 4s marks the start of the stationary phase, and the line at 12s marks the end of the stationary phase.

**Fig. 4** shows the peak foot positions averaged over perturbations during the moving and stationary phases relative to the start of the visual perturbation for all perturbation conditions. In MB, foot positions deviated significantly leftward in the moving phase (p < 0.001, *d* = - 1.663) and remained deviated to the left in the stationary phase (p < 0.001, *d* = 1.861). In both MT-HR and MT-EM, foot positions were not significantly different from zero in the moving phase (MT-HR: p = 0.271, *d* = 0.286; MT-EM: p = 0.827, *d* = -0.056). However, they significantly deviated to the right in the stationary phase (MT-HR: p = 0.018, *d* = 0.662; MT-EM: p = 0.039, *d* = 0.565). In MT-MR condition, during the stationary phase (0.024 ± 0.036 m), foot position deviations was 6 times larger than in the moving phase (0.006 ± 0.022 m). In MT-EM condition, the peak foot position was greater during the stationary phase (0.023 ± 0.041 m) than during the moving phase (−0.001 ± 0.021 m). We note that this effect was not very consistent between participants.

**Figure 4.**
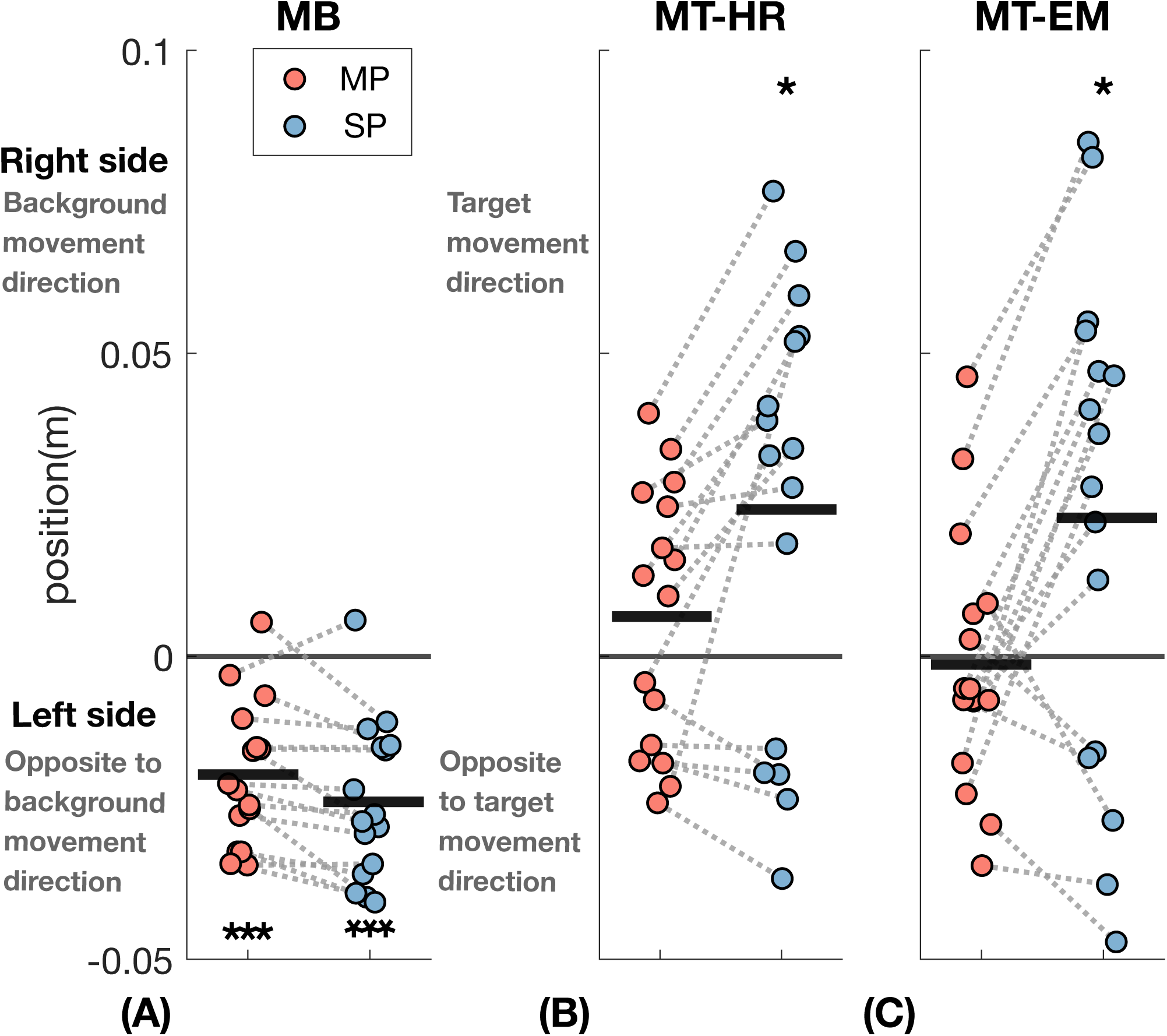
Averaged peak foot positions relative to the first sample of the visual perturbation epoch in the MP (moving phase) and SP (stationary phase), respectively. We averaged the two foot positions and selected the peak value during each phase during the visual perturbations. These peak values were then averaged across repeated visual perturbations as the mean (n = 16). The black thick lines represent the mean value across individuals. One-sample t-test was performed. Significant differences from zero are indicated by asterisks (* denotes p < 0.05, ** denotes p < 0.01, *** denotes p < 0.001). From left to right: (A) MB: moving background; (B) MT-HR: moving target with head rotation; (C) MT-EM: moving target with eye movement.

A two-way repeated measures ANOVA showed significant effects of Visual perturbation on averaged foot peak positions (F_visual perturbation_ (df = 1.390,20.854) = 17.006, p < 0.001, *η*^2^ = 0.380). Post-hoc results showed that MB was significantly different from MT-HR (p < 0.001, *d* = -1.392), and MT-EM (p < 0.01, *d* = -1.241). Phase also had a significant effect on averaged foot peak positions (F_phase_ (df = 1,15) = 9.406, p < 0.01, *η*^2^ = 0.055). The interaction between Visual perturbation and Phase (F_visual perturbation*phase_ (df = 2,30) = 8.869, p < 0.001, *η*^2^ = 0.053) indicated that during the stationary phase the differences in averaged foot peak positions between visual perturbations were larger than during the moving phase.

Qualitatively, in all conditions, the CoM appeared to follow the positions of the feet (**Fig. 5**). In MB, significant leftward deviations occurred in the moving phase (p < 0.001, *d* = -1.604), which remained significant in the stationary phase (p < 0.001, *d* = -2.135). In both MT-HR and MT-EM, peak CoM positions were not significantly different from zero in the moving phase (MT-HR: p = 0.401, *d* = 0.216; MT-EM: p = 0.983, *d* = -0.005). However, they deviated significantly rightward in the stationary phase (MT-HR: p = 0.026, *d* = 0.619; MT-EM: p = 0.041, *d* = 0.559). Generally, the peak CoM positions deviated leftward during MB, whereas in MT-HR and MT-ER, they deviated to the right.

**Figure 5.**
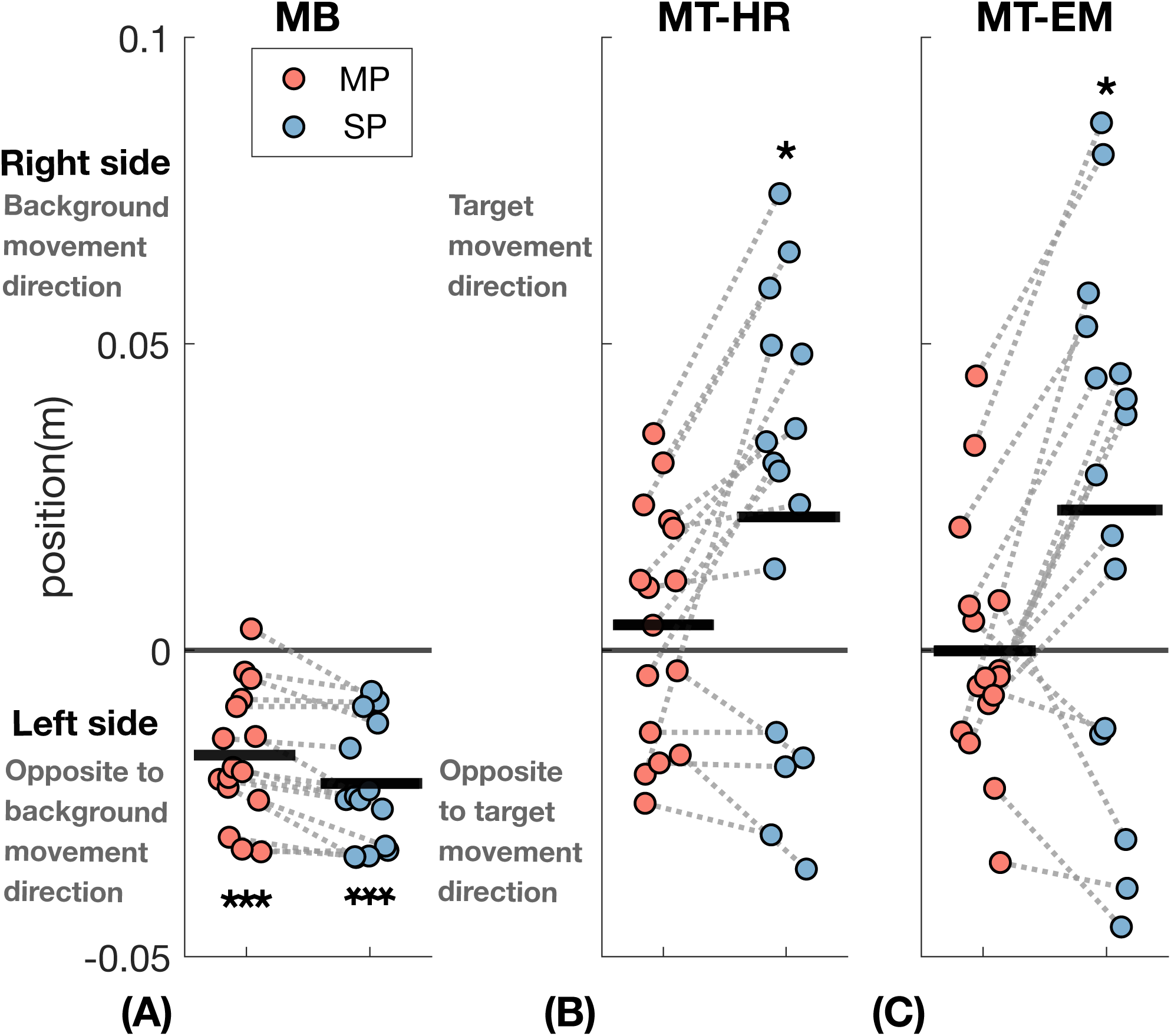
Peak CoM positions relative to the first sample of the visual perturbation epoch in the MP (moving phase) and SP (stationary phase). We selected the peak value in each phase during visual perturbations. These peak values were then averaged across repeated visual perturbations as the mean (n = 16). The black thick lines represent the mean value across individuals. One-sample t-test was performed. Significant differences from zero are indicated by asterisks (* denotes p < 0.05, ** denotes p < 0.01, *** denotes p < 0.001). From left to right: (A) MB: moving background; (B) MT-HR: moving target with head rotation; (C) MT-EM: moving target with eye movement.

Peak CoM positions in MB were significantly (F_visual perturbation_(df = 1.419, 21.278) = 14.918, p < 0.001, *η*^2^ = 0.347) smaller than in MT-HR (p < 0.01, *d* = -1.271) and MT-EM (p < 0.01, *d* = -1.209). While there was no significant difference between MT-HR and MT-EM. A significant effect of Phase was also observed (F_phase_(df = 1,15) = 8.273, p = 0.012, *η*^2^ = 0.056), showing that the CoM deviation in the stationary phase was larger than in the moving phase (p = 0.012, *d* = 0.470). The interaction between Visual perturbation and Phase (F_visual perturbation*phase_ (df = 2,30) = 9.061, p < 0.001, *η*^2^ = 0.056) indicated that during the stationary phase the differences in peak CoM positions between visual perturbations were larger than during the moving phase.

Peak frontal plane trunk orientations during the moving and stationary phases are shown in **Fig. 6**. In MB, trunk orientation did not significantly differ from zero in either phase (moving phase: p = 0.483, *d* = -0.180; stationary phase: p = 0.764, *d* = 0.077). In MT-HR, the trunk was significantly oriented leftward during both moving (p < 0.001, *d* = -1.713) and stationary phases (p < 0.001, *d* = -2.617). In MT-EM, trunk orientation was not significantly different from zero in the moving phase (p = 0.133, *d* = -0.397), while it deviated leftward in the stationary phase (p < 0.01, *d* = -0.912).

**Figure 6.**
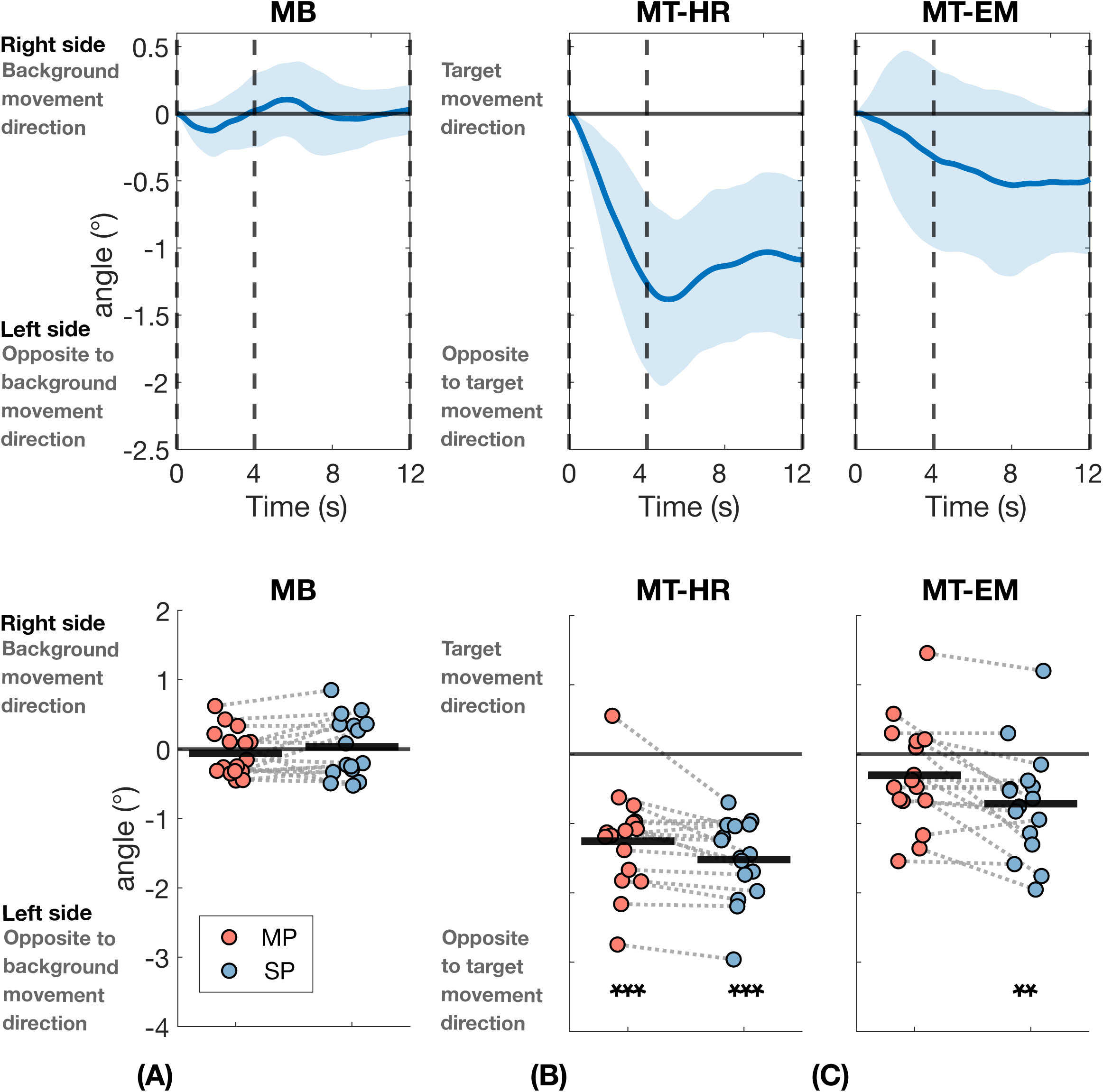
Deviations in frontal plane trunk orientation and peak trunk orientation during visual perturbation epochs. The panels in the first row show the deviations of trunk orientation. Data represent mean value across repeated visual perturbations and participants and the shaded area represents the between-participants standard deviations (mean ± s.d., n =16). The vertical dashed lines in the figures at the first row separate the different phases: the line at 0s marks the start of the moving phase, the line at 4s marks the start of the stationary phase, and the line at 12s marks the end of the stationary phase. The panels in second row show the peak frontal plane trunk orientation in MP (moving phase) and SP (stationary phase), respectively. We selected the peak value in each phase during visual perturbations. These peak values were then averaged across repeated visual perturbations as the mean (n = 16). The black thick lines represent the mean value across individuals. One-sample t-test was performed. Significant differences from zero are indicated by asterisks (* denotes p < 0.05, ** denotes p < 0.01, *** denotes p < 0.001). From left to right: (A) MB: moving background; (B) MT-HR: moving target with head rotation; (C) MT-EM: moving target with eye movement. All the data were referenced to the first sample of the visual perturbation epoch.

Peak trunk orientation depended on the visual perturbation condition (F_visual perturbation_(df = 2,30) = 24.581, p < 0.001, *η*^2^ = 0.559). In MT-HR, it was significantly smaller (more leftward) than in both MT-EM (p < 0.001, *d* = -1.395) and MB (p < 0.001, *d* = -2.183), but there was no significant difference between MB and MT-EM (p = 0.173, *d* = 0.789). A significant effect of Phase was observed as well (F_phase_(df = 1,15) = 16.271, p < 0.01, *η*^2^ = 0.016), indicating the leftward tilt became larger during the stationary phase. The interaction between Visual perturbation and Phase (F_visual perturbation*phase_ (df = 2,30) = 5.944, p < 0.01, *η*^2^ = 0.020) indicated that during the stationary phase, the differences in trunk orientation between visual perturbations were larger than in the moving phase.

The foot placement errors during the moving and stationary phases of each perturbation condition can be seen in **Fig. 7**. In MB, leftward (negative) foot placement errors were significant at the second (p < 0.01, *d* = -0.929), third (p < 0.01, *d* = -0.867), fourth (p < 0.01, *d* = -0.886), and sixth steps (p = 0.016, *d* = -0.679) during the moving phase. During the stationary phase, rightward (positive) foot placement errors were significant at the second (p = 0.035, *d* = 0.580), sixth (p < 0.01, *d* = 0.873), and eighth (p = 0.032, *d* = 0.593) steps. In MT-HR, rightward foot placement errors were significant at the fourth (p < 0.01, *d* = 0.769) and seventh (p < 0.01, *d* = 0.824) steps during the moving phase. During the stationary phase, rightward foot placement errors were significant at the first (p < 0.01, *d* = 1.011), second (p = 0.030, *d* = 0.601), third (p < 0.001, *d* = 1.107), fifth (p < 0.01, *d* = 0.838), sixth (p = 0.028, *d* = 0.610), seventh (p = 0.013, *d* = 0.702), ninth (p < 0.01, *d* = 0.806) and tenth (p = 0.024, *d* = 0.629) steps. In MT-EM, a leftward foot placement error was significant in the second step (p < 0.01, *d* = -0.817) during the moving phase. During the stationary phase, rightward foot placement errors were significant at the second (p = 0.021, *r_rb_* = 0.647), fifth (p < 0.001, *d* = 1.129), seventh (p < 0.01, *d* = 0.880), and ninth (p = 0.037, *d* = 0.573) steps.

**Figure 7.**
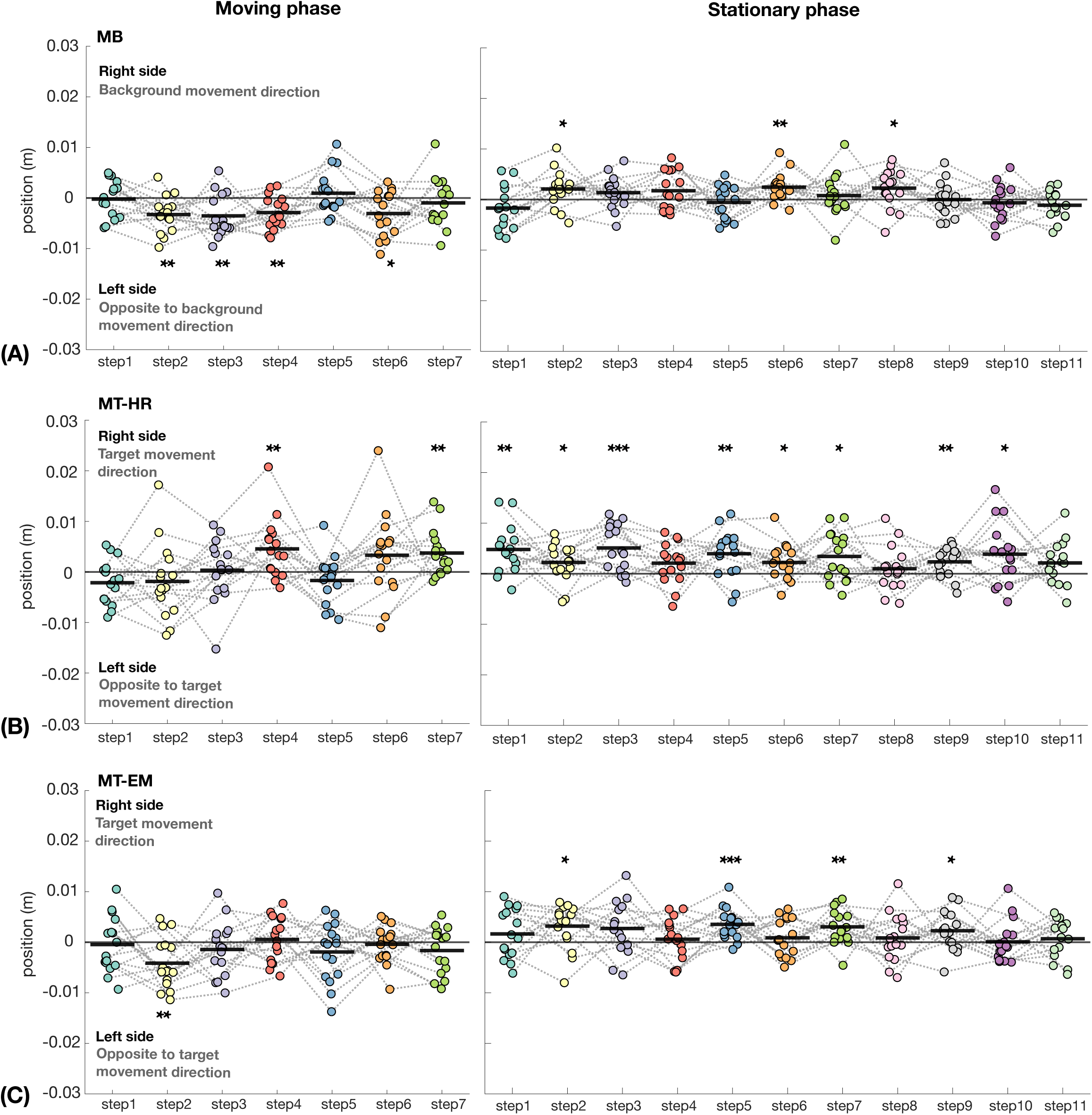
Foot placement errors in moving and stationary phases after the onset of the visual perturbation (n = 16). The black thick lines represent the mean value across individuals. One-sample t-tests were performed for each step. For steps which did not follow a normal distribution, we used Wilcoxon signed-rank tests instead. Steps with significant differences from zero are indicated by asterisks (* denotes p < 0.05, ** denotes p < 0.01, *** denotes p < 0.001). From top to bottom: (A) MB: moving background, (B) MT-HR: moving target with head rotation, (C) MT-EM: moving target with eye movement.

## Discussion

The visual system can be used to gather information from the environment to control balance. However, this information can in some situations be considered as a perturbation (Logan et al., 2010; Mohebbi et al., 2022). For example, on a stationary train, one may perceive to be moving when another train slowly passes by. In this study, we applied different types of visual perturbations to study if and how these affect balance during walking through eliciting changes in foot placement. Our results indicated that the variability of trunk sway and foot placement increased over the entire walking trial only when tracking a moving target, either with head rotation (MT-HR) or with eye movement only (MT-EM). During the visual perturbation epochs, fixating a stationary target while the background moved caused a deviation of foot and CoM trajectories opposite to the background movement. Tracking a moving target with head rotation or with eye movement only did not cause obvious deviations of foot and CoM trajectories during the moving phase, but did cause deviations in the target direction during the stationary phase.

Over the whole walking trial, a negative effect of visual perturbations on gait characteristics was only found in MT-HR and MT-EM. In these two conditions, step width variability, CoM position variability, trunk orientation variability, and RMS of foot placement errors were significantly higher than in NW. This indicates that visually tracking a moving target destabilizes walking even in healthy young participants. Unexpectedly, MB did not induce similar changes in gait characteristics except CoM variability. Although foot and CoM trajectories in MB deviated initially, they returned towards normal during the stationary phase. As a result, the effect of MB may have been diluted, which may have masked the overall impact of the perturbation on the trial.

During the visual perturbation epoch, the deviations in foot placement and CoM positions were different across conditions as well as different phases. In the current study, we used a 0.25 Hz cutoff frequency to filter the data and a validation at 4 Hz showed consistent results (**Fig. S1**). As expected, the deviation in foot placement during the moving phase varied between conditions. MB caused a deviation opposite to the background movement. However, in MT-HR and MT-EM, no obvious deviations were observed in this phase, in line with findings from a study by Vallis and Patla (2004). This may be explained by correction based on other information. For one, in the target tracking tasks, the CNS likely interpreted the relative background movement as a predictable outcome of self-motion, attenuating its perceptual salience and reducing the need for behavioral adjustments (Spering et al., 2011). Other sources of information that might help to distinguish self-motion and external motion may be: (1) the stationary background sensed by peripheral vision: the fixed black-and-white vertical stripes background provided a reference, (2) proprioception: proprioceptive information from the neck and eye extraocular muscles would indicate self-motion (Wang and Pan, 2013; Blumer et al., 2024), (3) the presence of an efference copy: predictions based on efference copy of the motor commands related to target tracking would also indicate self-motion (Wolpert and Ghahramani, 2000). In addition, since the onset of the target movements was unpredictable and fast, saccades may have been used during tracking instead of smooth pursuit movements (Leigh and Kennard, 2004). While smooth pursuit eye movements appear to disturb balance during walking, saccades were not found to do so (Thomas et al., 2017). Saccadic eye movements may induce suppression of visual information outlasting the duration of the saccade itself (Ibbotson and Krekelberg, 2011), which may have reduced sensitivity to the visual perturbation.

In the stationary phases, foot placement deviations were in opposite directions for the MB condition versus MT-HR and MT-EM conditions. In the MB condition, the deviations of the the foot trajectories subsequently decreased (**Fig. 3**). This suggests that the stationary target and background may have helped to correct the trajectory. The averaged peak foot placement (**Fig. 4**) and CoM position (**Fig. 5**) were slightly larger in the stationary phase, but this may have been caused by ongoing movement initiated during the moving phase and could be due to a shift in peak timing caused by the low-pass filtering. In the MT-HR and MT-EM conditions, the deviations of the foot trajectories reached significance only during the stationary phase and increased during this phase. Obviously, accumulation of small errors over the moving phase may have occurred, leading to significant deviations only in the stationary phase. This finding may also reflect that heading direction was aligned with gaze direction in this phase in all conditions. It has been proposed that head orientation or gaze direction provides the CNS with a frame of reference, and thus a change of this reference releases a steering synergy and affects heading direction (Hollands et al., 2002; Vallis and Patla, 2004; Bernardin et al., 2012; Belmonti et al., 2013). Finally, walking with a large head or eye rotation angle may be considered as a dual task, decreasing the control over heading, but it is not obvious why this would lead to deviations in a specific direction.

The deviation from average CoM position and velocity at midstance could explain 80% of variance in mediolateral foot placement (Wang and Srinivasan, 2014). We dubbed the remaining variance “foot placement errors”. Such foot placement errors could either be caused by the control itself or by perturbations. In current study, we expected foot placement errors during visual perturbation to point in the direction of the illusory self-motion. During the MB perturbations, where the background moved to the right, participants indeed stepped left of the predicted foot placement from the second to the fourth and sixth steps in the moving phase, which would align with an illusion of a deviation in the CoM state (position or velocity) in this phase. During the MT-HR perturbations, with a right target movement, as expected rightward foot placement errors appeared at the 4^th^ and 7^th^ steps. During the MT-EM perturbations, the second step (of the left foot) was unexpectedly placed leftwards. In the stationary phase, significant errors were in all conditions towards the right. In MB, rightward foot placement errors in the stationary phase coincided with the foot and CoM trajectories returning to the origin. While in the MT-HR and MT-EM conditions, rightward foot placement errors in the stationary phase coincided with a further rightward deviation of the foot and CoM trajectories. These results suggest that the stationary target location determined heading direction. In the MB condition, all sensory inputs would align with a forward heading. In MT-HR and MT-EM conditions, sensory information was likely conflicting, with tactile information from feet indicated a forward direction conflicting with gaze and in part neck orientation. The walking direction appeared to follow the information from the (neck and) head, indicating a top-down regulation.

Generally, a lateral step is associated with a force accelerating the CoM in the opposite direction (Reimann et al., 2018). However, none of the foot placement errors was associated with a subsequent movement of the CoM to the opposite side. Instead, in contrast with our expectation, the foot placement deviations coincided with a CoM displacement in the same direction (**Fig. S1**). While we expected more rightward foot placement to coincide with a more leftward acceleration of the CoM due to a higher leftward ground reaction force and vice versa, foot placement does not directly determine the horizontal ground reaction force exerted during the subsequent stance phase (van Dieën et al., 2025 preprint). One may interpret a more rightward foot placement to reflect an adaptation of the base of support to the ongoing CoM movement, which would fit with our data. In addition, CoM accelerations arising from the ground reaction force after foot placement may have been corrected through ankle moments bringing the CoM in the same direction (van Leeuwen et al., 2020). Our predictions were based on the expected role of visual information in sensing the state of the CoM and the role of foot placement in stabilizing the CoM trajectory, in the frontal plane. Alternatively, the conflicting findings may be attributable to the fact, already alluded to above, that the visual tasks may affect heading estimates and that foot placement also plays a role in control of heading.

A significant frontal trunk roll was found in MT-HR, which is consistent with previous findings (Patla et al., 1999; Vallis and Patla, 2004; van Andel et al., 2023). Van Andel et al. (2023) illustrated that head rotation was associated with frontal plane rotation of the trunk. Previous studies (Patla et al., 1999; Vallis and Patla, 2004) proposed that this frontal trunk roll is used as a strategy to push the CoM in the desired direction when making a turn and this would match with our participants walking in the direction of the target. However, in our study, trunk roll only occurred in MT-HR but not in MT-EM, where a similar deviation in gait direction was observed. Before making a turn, people usually rotate their head in the direction of the turn as well (Grasso et al., 1998; Hicheur et al., 2005; Dollack et al., 2019). Therefore, we assume that the trunk roll may be a result of a mechanical coupling between head rotation and trunk roll.

To further evaluate the plausibility of this mechanical coupling, we imposed the same visual perturbations in the same group of participants while standing. For each condition, five repeated standing trials were conducted. During recording, participants were instructed to fixate on the target, or track the target either by head or eye movement and stand still for 40s. In each trial, a visual perturbation was triggered once. The visual perturbation began around the 10^th^ second and lasted for 16s. The setting of the visual perturbations was the same as in walking. The presence of trunk roll when turning the head while standing (**Fig. 8**) supports the idea that this could result from a mechanical coupling.

**Figure 8.**
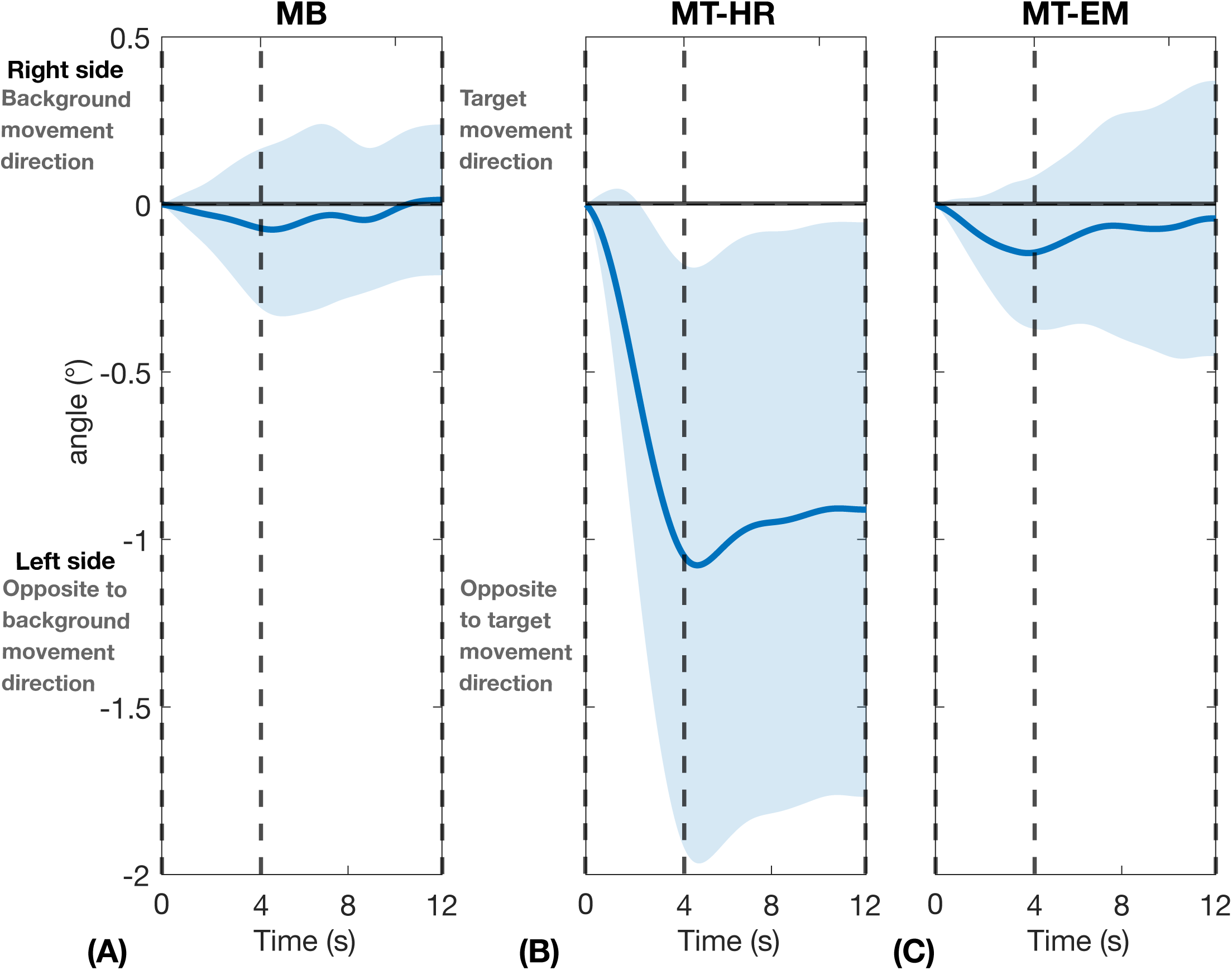
Frontal plane trunk orientation deviations during visual perturbation epoch in standing. Data represent mean value across repeated visual perturbations and participants and the shaded area represents the between-participants standard deviations (mean ± s.d., n =16). All the data were referenced to the first sample of the visual perturbation epoch. The vertical dashed lines in the figures separate the different phases: the line at 0s marks the start of the moving phase, the line at 4s marks the start of the stationary phase, and the line at 12s marks the end of the stationary phase. The shaded area represents the between-participants standard deviations. From left to right: (A) MB: moving background; (B) MT-HR: moving target with head rotation; (C) MT-EM: moving target with eye movement.

Although our experiment was conducted under controlled laboratory conditions, the observed interactions between gaze behavior and balance control likely reflect dual demands encountered in visually complex natural environments. In such settings, moving backgrounds and self-motion introduce visual perturbations that require integrated visual–vestibular–proprioceptive processing to maintain stability while tracking relevant targets. Many animal species integrate visual and other sensory inputs to support motor control, in ways that parallel human’s behaviour (Dauzere-Peres and Wystrach, 2024; Kodaka et al., 2004).

Evidence suggest that humans and non-human primates perceive visually induced self-motion through similar inferential mechanisms (Kodaka et al., 2004; Greenlee et al., 2016). The findings in the current study may therefore reflect general sensorimotor strategies that support effective locomotion and spatial orientation across species.

This study has several limitations. First, we only recruited healthy young adults, and results may be different, for example, in older adults, who have a high risk of falling (Lutz et al., 2008). Secondly, the experiment was conducted on a treadmill. This partially constrains walking speed and walking direction, which may have had an impact on participants’ responses to the visual perturbations. Thirdly, we did not assess foot orientation. Foot orientation may have played a role in redirecting the CoM trajectories. Further investigations may also consider the role of ankle moments for keeping balance during visual perturbations.

## Conclusion

Our results showed that visual perturbations challenge frontal plane balance, or in other words, the control of the CoM trajectories in this plane during walking. This was in line with expected effects on sensory estimates of the CoM state used for the control of foot placement in stabilizing the frontal plane CoM trajectory. Mechanical effects of head rotation, associated with visual tracking, had an additional disturbing effect on balance. However, effects of the gaze direction on heading appeared to affect gait simultaneously and cannot easily be distinguished from responses related to balance control.

## Acknowledgment

This study was supported by a scholarship (No. 202308310074) from the China Scholarship Council (CSC). The authors appreciate all participants for their voluntary participation. The authors thank Richard Casius, Leon Schutte and Paul Pesman for the technical assistance. The authors also thank Raven Huiberts and other collogues for their help in data collection.

## Competing interests

The authors declare no competing or financial interests.

## Data and resource availability

All data have been made available online. All files can be downloaded from: https://doi.org/10.5281/zenodo.17289747

## Notes

### Competing Interest Statement

The authors have declared no competing interest.

### Summary of Updates

In this version, we provided a description of a validation with a higher filter cutoff and have added actual numbers for key outcomes. In addition, we added a summary statement before the abstract and strengthened the discussion on the consistent direction between foot placement and CoM positions.

