## Supplemental Figure S1 and Figure S2 for "The Impact of Visual Perturbations on Balance Control during Walking"

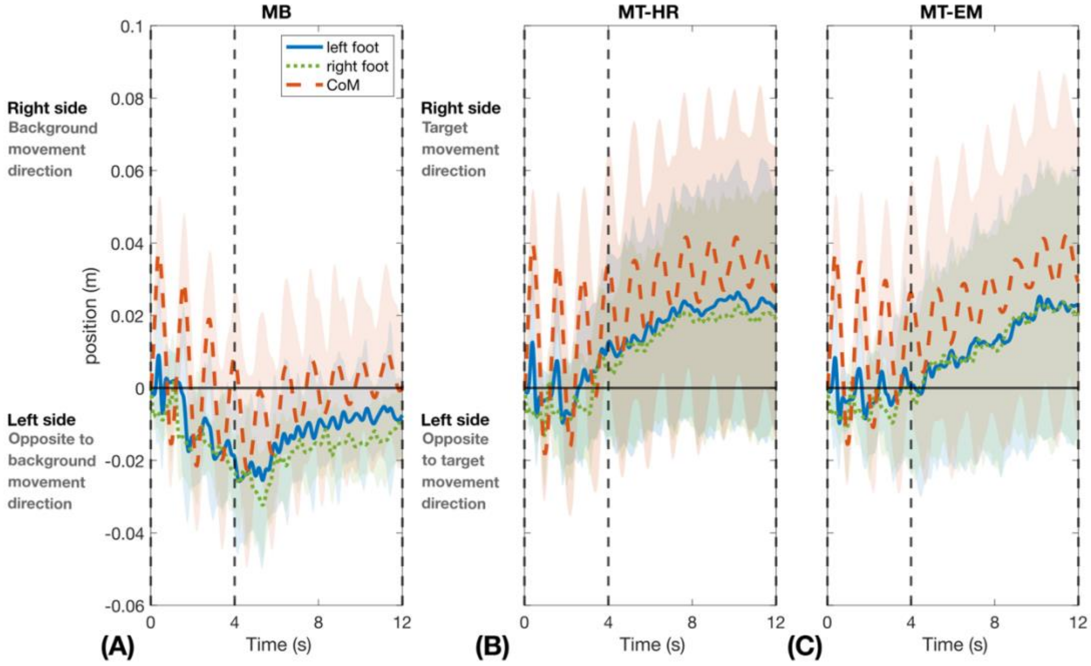


*Figure S1 Deviations in foot and CoM positions during visual perturbation epochs with a higher low pass filtered at 4Hz. Data represent mean value across repeated visual perturbations and participants and the shaded area represents the between-participants standard deviations (mean ± s.d., n =16). All the data were referenced to the first sample of the visual perturbation epoch. From left to right: (A) MB: moving background. (B) MT-HR: moving target with head rotation, (C) MT-EM: moving target with eye movement. The vertical dashed lines separate the different phases: the line at 0s marks the start of the moving phase, the line at 4s marks the start of the stationary phase, and the line at 12s marks the end of the stationary phase.*

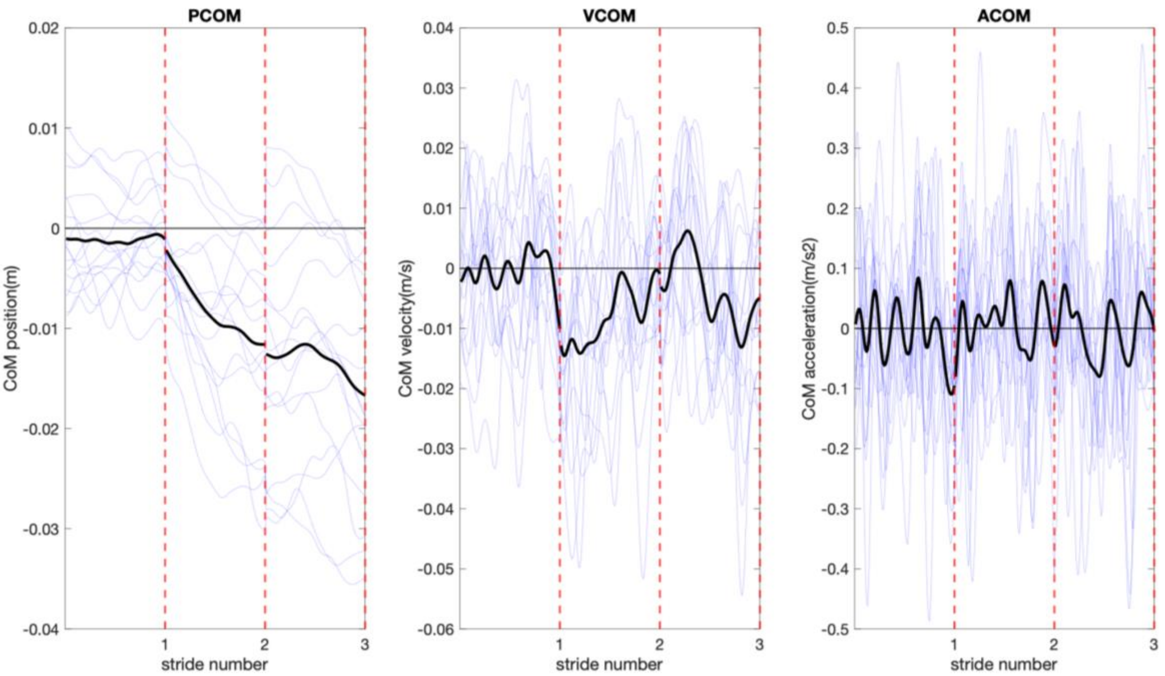


*Figure S2 CoM position, velocity, acceleration in the first three steps during moving phase in MB condition. The black curve in each panel represents averaged CoM position, velocity and acceleration across participants, respectively. The light blue lines represent individuals. The data were filtered at 4Hz over the whole time series and each stride was normalized to 100 samples. We averaged three unperturbed strides immediately preceding the perturbed steps. The first three strides after the onset of the perturbation were then extracted, and the averaged unperturbed strides were subtracted.*
